# Comparison of robotic automated and manual injection methods in zebrafish embryos for high throughput RNA silencing using CRISPR-CasRx

**DOI:** 10.1101/2023.07.04.547651

**Authors:** Joaquin Abugattas-Nuñez Del Prado, Yi Ding, Jan de Sonneville, Kees-Jan van der Kolk, Miguel A. Moreno-Mateos, Edward Málaga-Trillo, Herman P. Spaink

**Affiliations:** Institute of Biology Leiden, Animal Science and Health, Leiden University, Leiden, The Netherlands; Department of Biology, Universidad Peruana Cayetano Heredia, Lima, Perú; Life Science Methods BV, Leiden, The Netherlands; Andalusian Center for Developmental Biology (CABD), Pablo de Olavide University/CSIC/Junta de Andalucía, Seville, Spain; Department of Molecular Biology and Biochemical Engineering, Pablo de Olavide University, Seville, Spain

**Keywords:** CasRx, knockdown, zebrafish, high throughput, robot microinjector

## Abstract

Recently, the CRISPR-RfxCas13d (CasRx) system was proven to induce efficient mRNA knockdown in animal embryos. Here we compared the efficiency of CasRx-based RNA depletion with that of Cas9-mediated DNA targeting under the same conditions, using automated robotic and manual injection methods. As a proof-of-principle target, we used the no tail (tbxta) gene in zebrafish embryos, for which knockdown and knockout embryonic phenotypes were easy to be scored. Both Cas9 and CasRx systems induced loss of function phenotypes of *tbxta* gene. Higher percentage of severe phenotype was observed using Cas9 protein compared to the mRNA while the efficiency was similar in terms of Cas13d protein and mRNA. In addition, both the robotic and manual injection approaches yielded similar percentages of phenotypes and mortality rates. Therefore, our study not only showcases the potential of RNA-targeting CRISPR effectors for precise and potent gene knockdown, but also emphasizes automated microinjection in zebrafish embryos as an excellent alternative to manual methods for achieving gene knockdown at a high throughput level.

## INTRODUCTION

The zebrafish is a vertebrate model organism with many experimental advantages, such as large progeny, embryo transparency and amenability to genetic manipulation by microinjection. However, microinjection of the small zebrafish embryos is laborious and time-consuming. Therefore, better high throughput methods for the rapid microinjection of zebrafish embryos would significantly facilitate studies of gene function, pharmacological screens, and disease models.

Our laboratory previously showed that a robotic system allows for high throughput injections of CRISPR-Cas9 and DNA constructs into zebrafish zygotes with an efficiency comparable to that of manual injections [1]. Moreover, similar types of embryo injection robots were efficiently used for zebrafish tuberculosis infection studies and compound screens [2-5].

In addition to DNA modifications via CRISPR-Cas9 to generate F0 knockouts [6, 7], robotic injection also offers the possibility to generate and screen phenotypes resulting from various RNA depletion methods. For instance, RNA interference technologies and morpholinos offer an approach for RNA knockdown in several organisms. However, RNAi technologies have failed to be stablished in zebrafish model [8, 9] and morpholinos have shown toxicity, off-target effects, innate immune responses and discordant phenotypes with mutant animals [10-13].

Furthermore, CRISPR-Cas systems were firstly described as bacterial adaptive immune systems against phages [14-19]. They are organized in two main classes: Class 1 with Cas effectors composed of multiple subunits and Class 2 with monomeric large proteins; on top of that, specific subtypes within classes depends on the Cas endonuclease and its mechanism of action (Table 1). From Class 2, Cas9 and Cas12 are used for many genomic engineering applications [20-24] and have been applied in the zebrafish model [6, 25-27]. Moreover, the recently discovered Class 2 type VI Cas13 systems are RNA guided ribonucleases applied for transcriptomic engineering (Cas13a-Cas13d) [28-33]; in particular, the CRISPR-RfxCas13d (CasRx) system induces efficient knockdown of both zygotically and maternally provided RNAs in animal embryos [34].

**Table 1:** Classification of different CRISPR-Cas systems by classes and types known alongside with their number of subtypes, different Cas endonucleases and their target molecule [33]. “--” indicates that the mechanism of action and biological function is yet not confirmed for this type IV system.

| Class | Type | N. of Subtypes | Cas Endonuclease | Target |
| --- | --- | --- | --- | --- |
| Class 1 | Type I | 7 | Cas3 | DNA |
|  | Type III | 2 | Cas10 | DNA/RNA |
|  | Type IV | 2 | -- | -- |
| Class 2 | Type II | 3 | Cas9 | DNA |
|  | Type V | 7 | Cas12 | DNA |
|  | Type VI | 3 | Cas13 | RNA |

Here, we used automation to quantitatively evaluate the relative efficiencies of CRISPR-RfxCas13d and CRISPR-Cas9 to inactivate the function of the tbxta gene at the RNA or DNA level, respectively, and generate the resulting no-tail phenotype [35]. We show that high-speed robotic injection of the CRISPR-RfxCas13d achieves survival rates and knockdown efficiencies comparable with manual injections, thus opening the possibility of high throughput interrogation of the transcriptome in a tunable manner.

## RESULTS

### Establishment of robotic injection procedures

In this study we have developed and upgraded a robot based on previous versions [1, 3, 5]. The new version of the robot is a one-box fully integrated device with touchscreen and provides optimal injection speed into early-stage zebrafish embryos (Figure 1A). An agarose grid plate was prepared using a 9 × 100 well mold (Figure 1B). Zebrafish embryos can be placed in each grid well (Figure 1C). Subsequently, a needle was mounted into the needle holder, and needle tip (xy direction) and height (z direction) were calibrated (Figure 1D, video S1). For droplet calibration, a machine learning algorithm was added to recognize the shape of the droplet in a well filled with mineral oil and the volume of the droplet was given automatically (Figure 1E, video S2). Injections can be chosen either into middle of yolk (Figure 1F, video S3) or close to first cell (Figure 1F, video S4). Statistics on the top left of the screen give an overview of the number of embryos in each category while injection (Figure 1F, video S3 and video S4). Eggs where a first cell cannot be recognized (“no-cell” type) can be chosen to be injected or skipped. The skipped embryos are registered, and these eggs can be killed using a blunt needle (Figure 1G, video S5). After the injection and killing are finished, the embryos can be easily transferred to petri-dishes for culture (Figure 1H).

**Figure 1.**
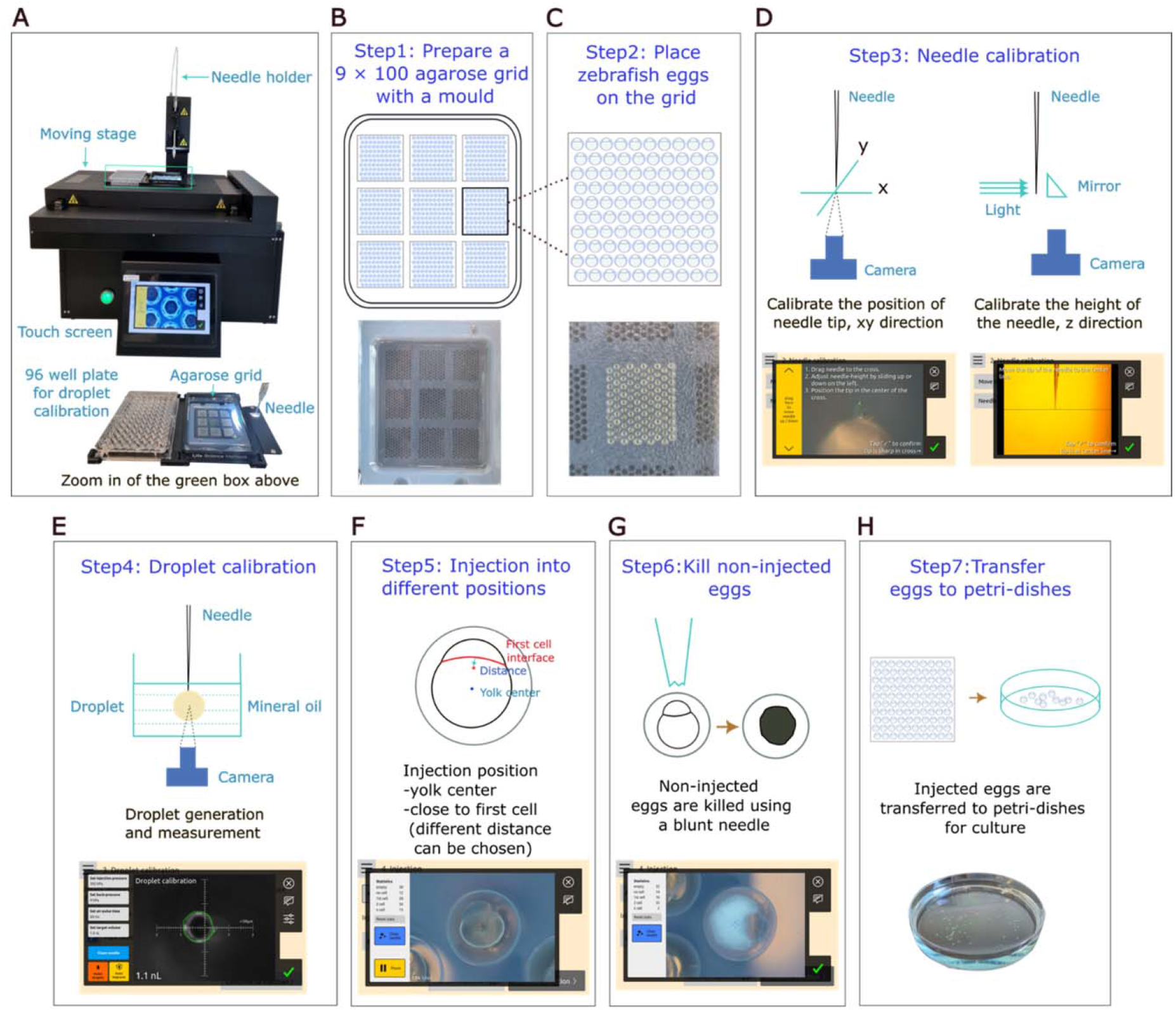
Overview of the zebrafish egg injection robot and experimental outline. **A.**A representative image of the integrate egg injector. Needle holder is used to load needles and motorized stage enables the movement of plates containing embryos during injections. 96 well plate is for droplet calibration and the agarose gel grid plate is for placement of zebrafish eggs. Touch screen allows the operation and visualization of all the injection processes. B-H. The procedure of zebrafish embryos injection using the robot. **B**. Step1: Preparation of an agarose grid plate using a mold. Different molds e.g., 9 × 100, 1024, 360 grid wells, can be used based on research purposes. **C**. Step2: Placement of zebrafish eggs on the agarose grid. Zebrafish embryos can b placed randomly at any wells of the grid as empty wells can be recognized and skipped by the robot. **D**. Step3: Needle calibration for both xy direction and z direction as shown on the screenshots from the touch screen. **E**. Step4: Droplet calibration is performed by injection of suspension into mineral oil and droplet size can be determined automatically by the robot. **F**. Step5: Injection positions should be chosen either yolk center or close to first cell, subsequently the robot can do injections automatically for wells that users selected. Different injection distance, e.g., 0μm, 30μm and 50μm, can be chosen for close to first cell injection. **G**. Step 6: Non-injected eggs can be killed by loading a blunt needle. **H**. Step 7: After the injection is finished, the injected zebrafish eggs can be transferred to petri-dishes for culture and dead eggs should be removed.

### Comparison of manual and automated robotic injection

To compare injection results between manual and robotic injections, both CRISPR-Cas9 and CRISPR-RfxCas13d systems targeting one gene tbxta were used. Knockdown of the *tbxta* gene using both the two systems resulted in different level of tail and notochord defect phenotypes at 30 hpi (Figure 2A). The CasRx gRNA used to reproduce this phenotype was previously validated by RT-qPCR, resulting in an average transcript level decrease of 60% (34). Additionally, the Cas9 gRNAs achieved more than 90% of F0 biallelic knockouts, which were validated by DNA sequencing (6). Higher percentage of embryos with severe phenotype (type III) and lower percentage with mild phenotypes (type I and type II) were observed in Cas9 protein than Cas9 mRNA groups (Figure 2B). The percentage of embryos with different types of phenotypes between Cas13d protein and Cas13d mRNA groups showed no distinct differences (Figure 2C). Furthermore, the knockdown efficiency of both Cas9 and CasRx using a robot was highly comparable with manual injector (Figure 2B and 2C). Injection into the cell interface and yolk center of zebrafish embryos showed no difference in the phenotype (Figure 2B and 2C). The mortality rate of injected embryos was similar between manual and robotic injection (Table 2). However, the speed of the robot (59 ± 3 embryos/minute, 2169 injected embryos) was 3 times faster compared to the manual injections (20 ± 3 embryos/minute, 990 injected embryos) (Table 2).

**Table 2:** **Different characteristics of manual and robot injection methods**. Mortality rate 6hpi, 30hpi are expressed in percentage ± STD. Injection speed is expressed in injected embryos per minute ± STD. Total injected embryos is the number of injected embryos. At least 3 independent experiments were performed.

|  | Manual | Robot |
| --- | --- | --- |
| Mortality rate 6hpi | 3.9 ± 0.8 | 3.3 ± 1.1 |
| Mortality rate 30hpi | 3.8 ± 1.5 | 4.7 ± 1.3 |
| Injection speed (embryos/minute) | 19.7 ± 3.1 | 58.9 ± 2.6 |
| Total injected embryos | 1731 | 3603 |

**Figure 2:**
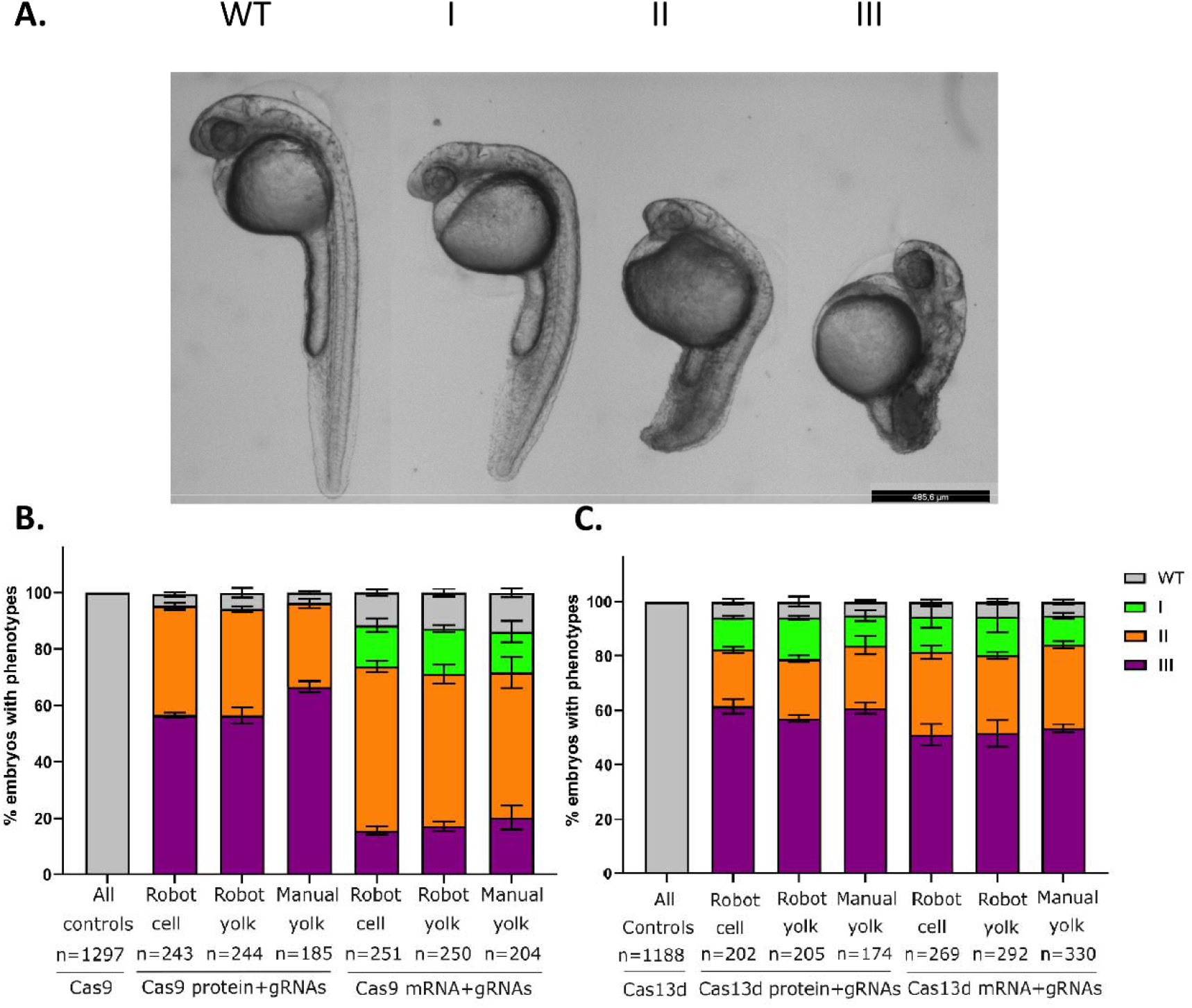
Percentage of zebrafish embryos with different grades of no tail phenotype caused by *tbxta* gene knockdown. **A**.Knockdown of the *tbxta* gene using both Cas9 and CasRx systems results in different levels of no tail phenotype. Representative phenotype images are acquired from CasRx injections. WT, normal tail and notochord; type I, short tail and normal notochord; type II, notochord development failure and short tail; type III, notochord development failure and shorter tail. Scale bar 485.6μm. **B**. Percentage of embryos of each grade of no tail phenotype caused by Cas9 protein or mRNA targeting *tbxta* gene using automatic robot or manual injections. As all the control groups injected either with Cas9 protein or mRNA only show wild type phenotype, therefore we combined all the control groups to one bar, namely “All controls” (Cas9 protein robot, n=434; Cas 9 protein manual, n=213; Cas9 mRNA robot, n=447; Cas9 mRNA manual, n=203). **C**. Percentage of embryos of each grade of no tail phenotype caused by CasRx protein or mRNA targeting tbxta gene using automatic robot or manual injections. As all the control groups injected either with Cas13d protein or mRNA only show wild type phenotype, therefore we combined all the control groups to one bar, namely “All controls” (Cas13d protein robot, n=334; Cas 13d protein manual, n=173; Cas13d mRNA robot, n=432; Cas13d mRNA manual, n=249).

## MATERIALS AND METHODS

### Zebrafish maintenance and embryo production

Leiden University researchers follow the European rules for the use of animals. Zebrafish wildtype strains ABTL are maintained and bred according to standard conditions [36]. One cell stage ABTL embryos were used in this study.

### CasRx tbxta gRNA production

The gRNAs for CasRx targeting tbxta [34, 37] gene were produced by fill-in PCR with Phire HotStart II DNA polymerase (ThermoFisher Scientific, F122S) (primers in Sup. Table 1, IDT) in a thermal cycler (BioRad, T100) with 10 uM final concentration of each primer and the following program: 3 min at 95 °C, 35 cycles of 30 sec at 95°C, 30 sec at 60°C, and 30 sec at 72°C, and a final step at 72°C for 5 min. The PCR product was in vitro transcribed (Epicentre, ASF3507) under the control of promoter T7 for 12 hours at 37°C and DNAse treated with TURBO-DNAse for 20 min at 37°C. Finally precipitated with 3 M ammonium acetate and resuspended in nuclease-free water. gRNA integrity was verified with electrophoresis (Sup Fig 3), quantified by Nanodrop 2000 (Thermo Scientific) and titrated until high phenotypical penetrance was obtained. We observe that quantifying gRNAs after in vitro transcription can yield inaccuracies, which may lead to an overestimation of the amount of gRNAs being utilized. However, we mitigated this issue by titration of the gRNA employed.

### Cas9 and CasRx mRNA production

Plasmids containing Cas9 and CasRx genes, pT3Ts-nCas9n (Addgene 46757) and pT3Ts-RfxCas13d-HA (Addgene 141320) respectively were linearized with XbaI (NEB R0145L) for 3 hours at 37°C and then in vitro transcribed under control of promoter T3 (Ambion AM1348) for 3 hours at 37°C and DNAse treated with TURBO-DNAse for 20 min at 37°C. Capped mRNA from these reactions was purified with RNAeasy MiniKit (Qiagen 74104). RNA integrity was verified by electrophoresis (Sup Fig 1, 2) and quantified with Nanodrop 2000 (Thermo Scientific).

### CasRx protein expression and purification

Competent Rosetta DE3 pLysS (Novagen 70956) were transformed with the plasmid pET28b-RfxCas13d-His (Addgene 141322), which contain the recombinant CasRx gene. Cells were grown for 3 h at 37°C and then induced with 0.1mM IPTG for another 3 h. Cells were washed once with 20mM Tris-HCl pH 7.5 and the pellet was frozen at -80°C before next steps.

Pellets were resuspended with Lysis buffer (50 mM HEPES*KOH pH 7.5, 500 mM KCl, 10% v/v glycerol, 1mM DTT and 10 mM imidazole) and sonicated (QSonica, Q125) using the following program: 3 sec ON, 10 sec OFF, 120 cycles, 30% amplitude. Lysate was centrifuged and then filtered using a 0.2 um cellulose acetate filter. CasRx protein was then purified using an HisPur™ Ni-NTA spin column (Thermo Scientific, 88226) and washed with 20 column volumes of lysis buffer by gravity. Elution was performed in five steps with Elution buffer (50 mM HEPES*KOH pH 7.5, 500 mM KCl, 10% v/v glycerol, 1 mM DTT and 500 mM imidazole) diluted with Lysis buffer to get increasing concentrations of imidazole (10 mM, 30 mM, 50 mM, 100 mM and 200 mM). Fractions containing purified CasRx were pooled and then concentrated and dialyzed with Dialysis buffer (50 mM HEPES-KOH pH 7.5, 250 mM KCl, 1 mM DTT and 10% glycerol) using a 50K Amicon Ultra-15 (Millipore UFC905024) until concentration of 3 ug/ml was reached. Protein concentration was estimated with Nanodrop 2000 (Thermo Scientific) and extraction quality was verified by SDS-PAGE (Sup Fig 4). Single use aliquots of 4 ul were stored at -80°C.

### Cas9 protein and gRNAs

Alt-R™ – Cas9 protein, Alt-R™ crRNA and Alt-R™ tracrRNA were purchased from Integrated DNA Technologies (IDT). The same gRNAs (Sup Tab 1) were used in both mRNA and protein injections of Cas9. Alt-R CRISPR systems from IDT are optimized genome editing tool with predesigned gRNAs, to produce on-target double-strand breaks using high purity solutions.

Production of mRNAs, gRNA and protein CasRx and Cas9 (see methods) showed a high rate of capped mRNA production from linearized DNA template (∼100-fold ng of RNA α ng of DNA, data not shown), with little degradation after purifying as shown in the agarose gels (Sup Fig 1, 2). Similarly, the gRNAs generation showed higher efficiency (∼800-fold ng of RNA α ng of DNA, data not shown), having a reaction time 4 times longer than mRNA reaction and with low degradation (Sup Fig 3). The CasRx protein production was induced and purified with a Ni-NTA column followed by concentration with an Amicon ultracentrifugation column. We obtained a concentration of 3 µg/ml with a purity off approximately 85% (Sup Fig 4).

### Microinjection solutions and injection quantities

Cas9 gRNAs were prepared by annealing crRNA with tracrRNA using Duplex Buffer (IDT) and incubating at 95°C for 5 min. 3 different gRNAs were pooled and then mixed with Cas9 mRNA. Final quantity injected with 1 nl per embryo was 200pg of Cas9 mRNA and 1070 pg of gRNAs (357 pg each). Control groups were only injected with 200pg Cas9 mRNA. To produce the Cas9 ribonucleoprotein (RNP), the same gRNAs are mixed with Cas9 protein (IDT) and incubated at 37°C for 5 min. Final quantity injected with 1 nl per embryo is 5029 pg of Alt-R Cas9 protein and 1070 pg of gRNAs (357pg each). Control groups were only injected with 5029pg Cas9 protein. In the case of CasRx, the mRNA was mixed with the gRNA targeting tbxta transcript. Final quantity injected with 1 nl per embryo is 200 pg of mRNA and 300 pg of gRNA. Control groups were only injected with 200pg CasRx mRNA. CasRx protein was mixed with gRNA targeting tbxta transcript. The final quantity injected with 1nl per embryo is 3 ng of protein with 300 pg of gRNA. Control groups were only injected with 3ng CasRx protein.

### Manual microinjection

Manual microinjection was performed using a Pneumatic Pico-Pump (PV 820, WPI) and all embryos were injected in the yolk during one cell stage with around 5-8 psi depending on the needle pore size. All experiments were performed with at least three replicates.

### Automated robot microinjection and software updates

The detailed procedure of using the upgraded automated microinjection system (Life Science Methods. B.V. the Netherlands) was described in Figure 1. Some software updates have been carried out to improve the automated injection throughput and efficiency. A deep-learning network was updated to recognize the yolk center and the first cell interface. Another deep learning network has been added to recognize the shape of the droplet and compute to volume. The speed is increased from 1.8 second per injection in the previous version to 1.0 second per injection on average in the current system.

## DISCUSSION

In this study, we present a detailed procedure for utilizing an advanced zebrafish embryo injection robot and conduct a comparative analysis of injection outcomes between robotic and manual methods. The robotic system offered several advantages. The implementation of this robot allows individuals without any prior zebrafish egg injection experience to perform large-scale injections effortlessly. Additionally, the inclusion of a user-friendly touch screen interface enhances the operational convenience and provides a visually appealing display of the injection processes, ensuring a comfortable experience for the operator. Moreover, the robot offers the advantage of precise injection volumes by automatically calculating droplet sizes using a 2D image analysis. This automated computation significantly reduces errors associated with manual estimation, minimizing the potential translation of such errors into inaccuracies in injected volume. Lastly, the robotic system incorporated a feature where it recorded the positions of non-injected eggs and subsequently performed selective euthanasia, ensuring that only the injected embryos remained alive for further analysis.

Our results indicate that both the robotic and manual injection approaches yielded similar percentages of phenotypes and mortality rates, specifically regarding the knockdown of the *tbxta* gene. However, it is noteworthy that the robotic method exhibited superior speed, being three times faster than manual injections. This improvement in speed is attributed to the expanded region of interest, which allows for simultaneous evaluation of the next well during the ongoing injection process. On average, the robot achieved an injection speed of 1.0 second per injection, as demonstrated in video S3 and S4.

In the context of gene editing, traditional practices involved injecting DNA/mRNAs/gRNAs into the first cell of zebrafish embryos. However, recent studies, such as the work by Kroll et al. (6), have demonstrated that injection directly into the yolk is adequate for achieving F0 knockouts using the highly efficient CRISPR-Cas9 technique. In our study, we utilized a combination of multiple gRNAs targeting the *tbxta* gene and observed that yolk injections resulted in a substantial occurrence of tail and notochord defects. Moreover, we found that the proportion of phenotypes caused by yolk injection closely resembled those resulting from cell interface injection using the robotic system. It is important to note that our injections were administered at the one-cell stage, shortly after fertilization, minimizing the time elapsed before the cell started to inflate. Consequently, we propose that the timing of injection, specifically immediately following fertilization and prior to cellular expansion, holds greater significance than injecting into the zygotic cell itself [6].

It has been observed that Cas9 protein injections lead to more pronounced phenotypic effects compared to Cas9 mRNA injections in terms of tbxta gene knockout. This disparity may be attributed to the ability of Cas9 protein to perform DNA double strand breaks (DSB) from 1-cell stage at the moment of injection; on the other hand, the mRNA would have to be translated for the protein to cause a DNA DSB. Moreover, our study also highlighted the potential of the CasRx system at the RNA level. The CasRx system for knockdown has undergone significant improvements through upgraded protocols and platforms in recent years [37]. In the present study, we utilized previously validated CasRx gRNAs targeting tbxta mRNA [34, 37], taking advantage of the enhanced CasRx system. Given that CasRx protein is not commercially available, we expressed and purified the protein in our own laboratory. Both the purified Cas13d protein and Cas13d mRNA injections resulted in high phenotypic penetrance. The percentage of phenotypes observed in the Cas13d protein group is similar with that in the mRNA group. This low difference between mRNA and protein injection of CasRx may be because our target gene tbxta is involved in embryo notochord formation, which occurs relatively late in zebrafish development. Nevertheless, if an early development gene is studied, the CasRx protein might have a higher phenotypic penetrance. Moreover, after the CasRx mRNA injection, protein translation in the embryo has not yet been studied in different developmental stages and the phenotypic penetrance would also depend on the target mRNA levels in these stages. Another interesting observation is that we can see Grade I phenotypes with the CasRx experiments. This could be explained by the previous validation for the used gRNA, which showed a reduction of mRNA levels of about 60% of the tbxta transcript by RT-qPCR (34, 37), as we expect the same efficiency in our experiments. This reduction would still allow for a low expression of this gene in the embryo. Additionally, it was demonstrated that disrupting a single nucleotide in the tail of the gRNA (CasRx binding domain) and using a 23-nucleotide binding sequence significantly improves the phenotypic penetrance (39). Specifically, the percentage of grade III no tail cases doubles from 35% to 70% with this chemically synthetized highly efficient gRNA structure (37).

Overall, our study contributes to the understanding of the benefits and feasibility of robotic automated injection methods in zebrafish embryos for gene silencing experiments. The combination of precise injection control and increased efficiency provided by the robotic system holds promise for accelerating research in early embryonic development and other areas that require large-scale injections. Furthermore, the exploration of the CasRx system showcases the potential of RNA-targeting CRISPR effectors for precise and potent gene knockdown at high throughput.

## SUPPLEMENTARY INFORMATION

Additional file 1: Video S1. Needle calibration by the robot. In the needle calibration interface, click “Move stage to mount needle”, the robot will move the needle holder up and the stage to a back position for mounting a needle. After a filled needle is mounted, the robot automatically brings the needle through a hole in the stage and a screen shows calibration for the xy direction of the needle tip. Drag the yellow area up or down until a pointed dot is shown and put the dot in the center of the red cross and tap ⍰ afterwards. Then, the robot brings the needle a bit lower, and positions the stage such that the needle is close to a prism mounted in the back-wall of the hole, to view the needle from the side. In the next screen, move the needle tip to the line and tap ⍰. The needle is calibrated.

Additional file 2: Video S2. Droplet calibration by the robot. In the droplet calibration interface, click the icon next to “Calibrate using a well plate”, select the well positions of the 96-well plate that are filled with mineral oil and water. Make a droplet by clicking “Make droplet” button and the volume of the droplet will be shown on the screen. Different injection settings can be adjusted to achieve a proper volume. “Auto improve” button aims to reach the target volume by adjusting injection pressure and air-pulse time automatically. “Clean needle” can be used when the needle is clogged.

Additional file 3: Video S3. Automatic injection into middle of yolk by the robot. In the injection interface, click “middle of yolk” and different type of embryos can be chosen for injection. Select the areas where embryos are placed and click “start injection”. The robot scans the agarose grid and perform injections into the yolk center of selected type of embryos. Statistics show the number of embryos in each category: empty, empty wells without embryos; no cell, embryos with no cell visible; 1_st_ cell, embryos with one cell; 2 cell, embryos with two cells; n cell, embryos with more than two cells. ETA shows the time left for injections.

Additional file 4: Video S4. Automatic injection into close to first cell by the robot. In the injection interface, click “Close to first cell”, the injection site can be chosen at a user specified distance from middle of the cell interface towards the yolk center as the positive direction. If “Inject even when no cell is visible” is chosen, the robot will inject a random position of those embryos. Select the areas where embryos are placed and click “start injection”. The robot scans the agarose grid and perform injections into close to first cell of embryos. Statistics show the number of embryos in each category: empty, empty wells without embryos; no cell, embryos with no cell visible; 1 _st_cell, embryos with one cell; 2 cell, embryos with two cells; n cell, embryos with more than two cells. ETA shows the time left for injections.

Additional file 5: Video S5. Kill non-injected embryos by the robot. If there are embryos without injection, click “Kill non-injected eggs” when the injection is finished and mount a blunt needle and perform the needle calibration as before. The robot remembers the positions of non-injected embryos and perform killing.

Additional file 6: Figure S1. Cas9 mRNA production. Electrophoresis of DNA and denatured RNA in an ethidium bromide stained 1.5% agarose gel. L. Smart Ladder (MW-1700-10, Eurogentec). 1. pT3Ts-nCas9n plasmid. 2. pT3Ts-nCas9n plasmid digested with XbaI. 3. Cas9 denatured capped mRNA.

Additional file 7: Figure S2. CasRx mRNA production. Electrophoresis of DNA and denatured RNA in an ethidium bromide stained 1.5% agarose gel. L. Smart Ladder (MW-1700-10, Eurogentec). 1. pT3TS-RfxCas13d-HAplasmid. 2. pT3TS-RfxCas13d-HAplasmid digested with XbaI. 3. CasRx denatured capped mRNA.

Additional file 8: Figure S3. CasRx gRNA for tbxta. Electrophoresis of denatured RNA in an ethidium bromide stained 2% agarose gel. L. Smart Ladder (MW-1700-10, Eurogentec). 1. Denatured gRNA for targeting tbxta gene.

Additional file 9: Figure S4. CasRx protein production. SDS-PAGE of different samples during the CasRx purification process with E. coli Rossetta DE3 pLyss transformed with pET28b-RfxCas13d-His. Red arrows show the CasRx protein. L. ladder mPAGE Unstained Protein Standard (MPSTD3, Millipore). 1. Pre induction sample. 2. Post induction sample (IPTG 0.1mM). 3. Input reference after sonication. 4. Input reference after filtration with 0.2um cellulose acetate filter. 5. Flowthrough reference after the Ni-NTA column. 6. Wash reference. 7. Elution 1. 8. Elution 2. 9. Elution 3. 10. Elution 4. 11. Elution 5. 12. Elution 6.

Additional file 10: Table S1. Primers and RNA sequences

## ABBREVIATIONS

CasRx: RfxCas13d
gRNA: guide RNA
mRNA: messenger RNA
crRNA: CRISPR RNA
tracrRNA: trans-activating CRISPR
RNA RNP: ribonucleoprotein
Hpi: hours post injection

## AKNOWLEGDMENTS

We would like to thank Guus van der Velden, Ulrike Nehrdich and Natasha Montiadi for helping us taking care of zebrafish. We are also grateful for B.E.V. Koch, PhD help for discussion and suggestions during CasRx protein production standardization and helping with manuscript submission.

## AUTHOR CONTRIBUTIONS

JANDP expressed and purified CasRx protein, produced CasRx and Cas9 mRNA and CasRx gRNAs. JANDP an YD performed the injection experiments and wrote the first version of the manuscript. JdS designed the robotic injection equipment, K-JvdK designed the software for robot control and machine learning, JdS, EMT, MAM-M and HPS supervised the study and provided materials. All authors have contributed to writing this article.

## DATA AVAILABILITY

All raw data is available on request

## ETHICS APPROVAL AND CONSENT TO PARTICIPATE

Not applicable.

## CONSENT FOR PUBLICATION

Not applicable.

---

**Additional file 6: Figure S1.**
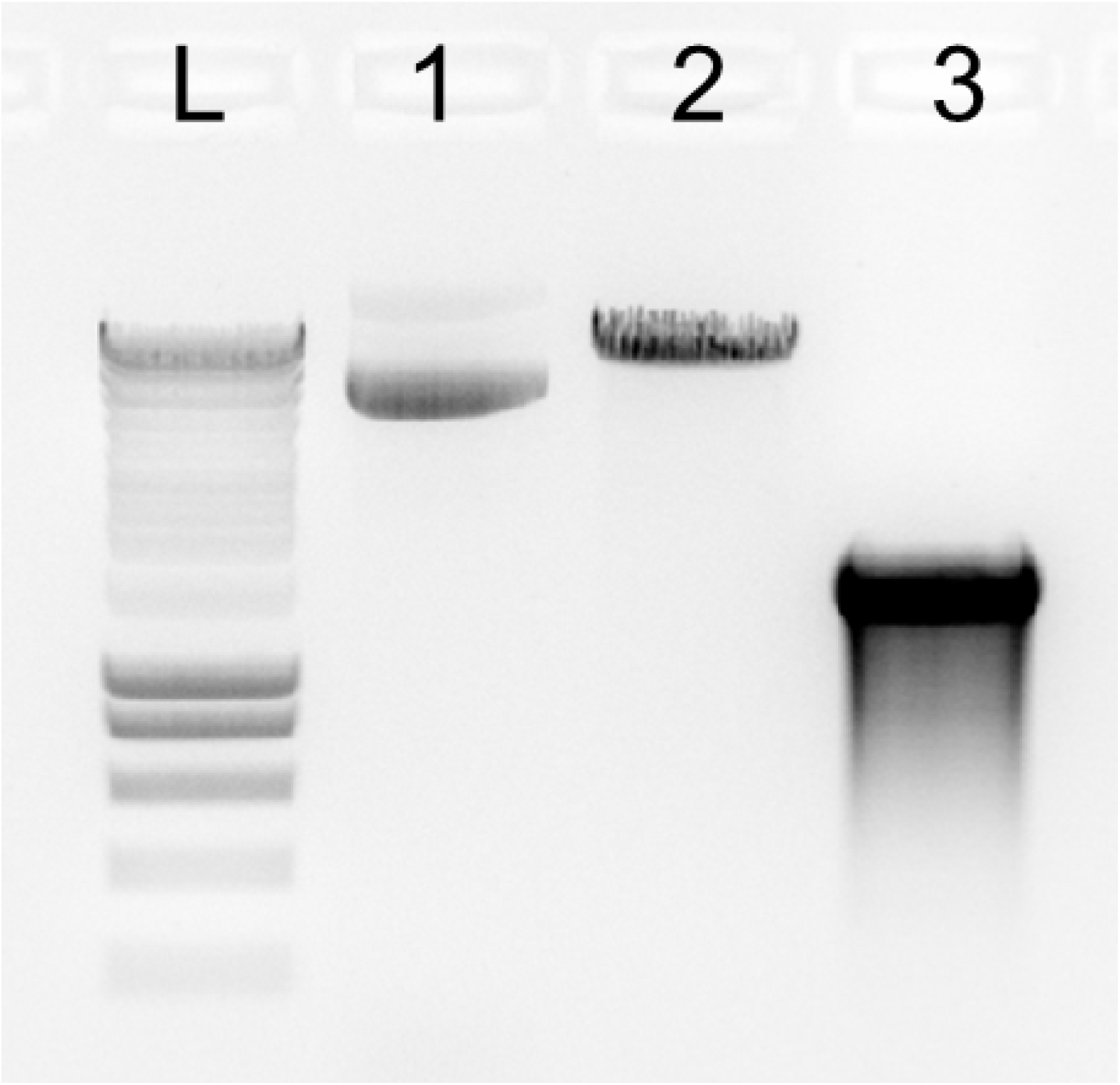
Cas9 mRNA production.

**Additional file 7: Figure S2.**
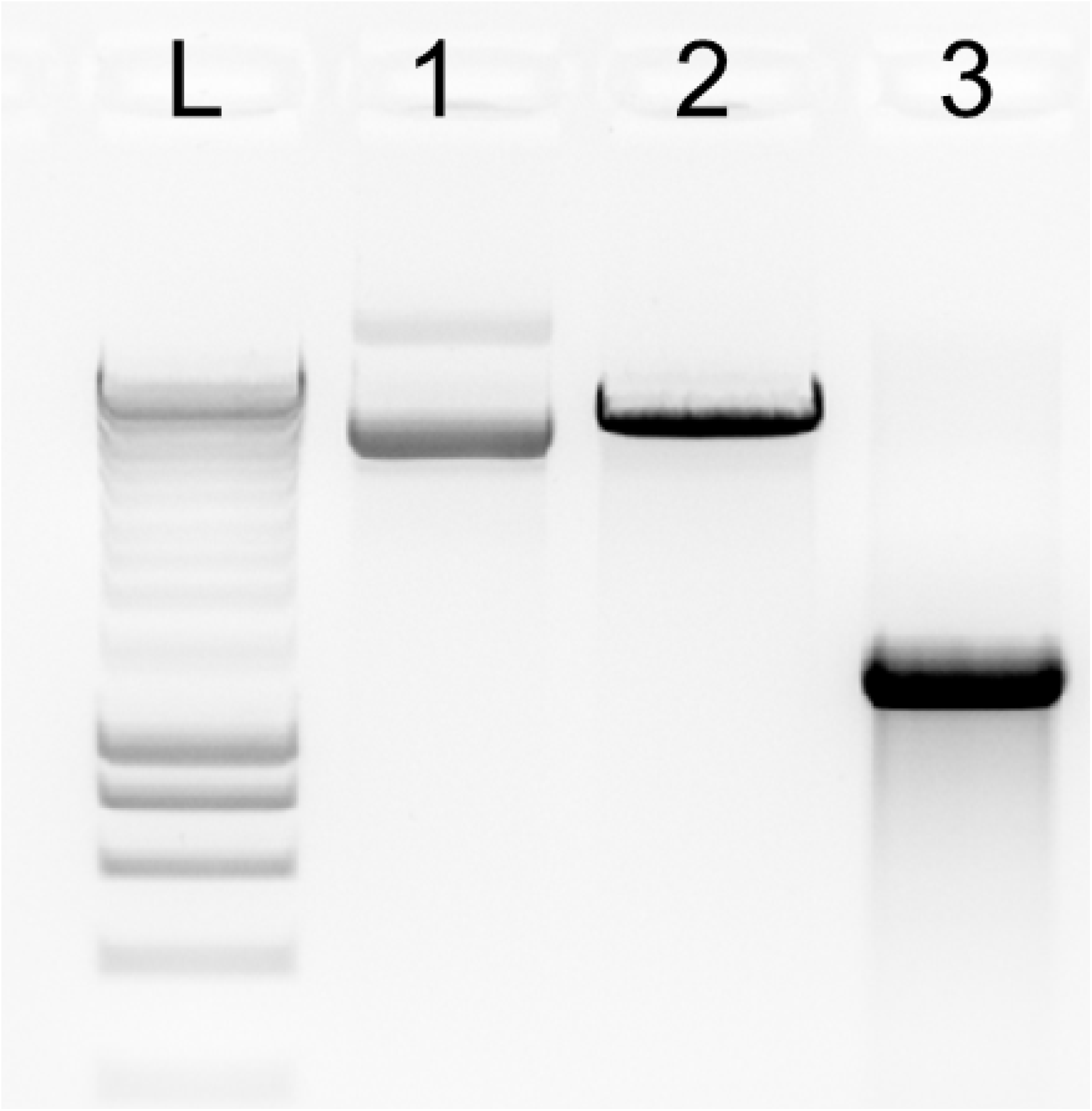
CasRx mRNA production.

**Additional file 8: Figure S3.**
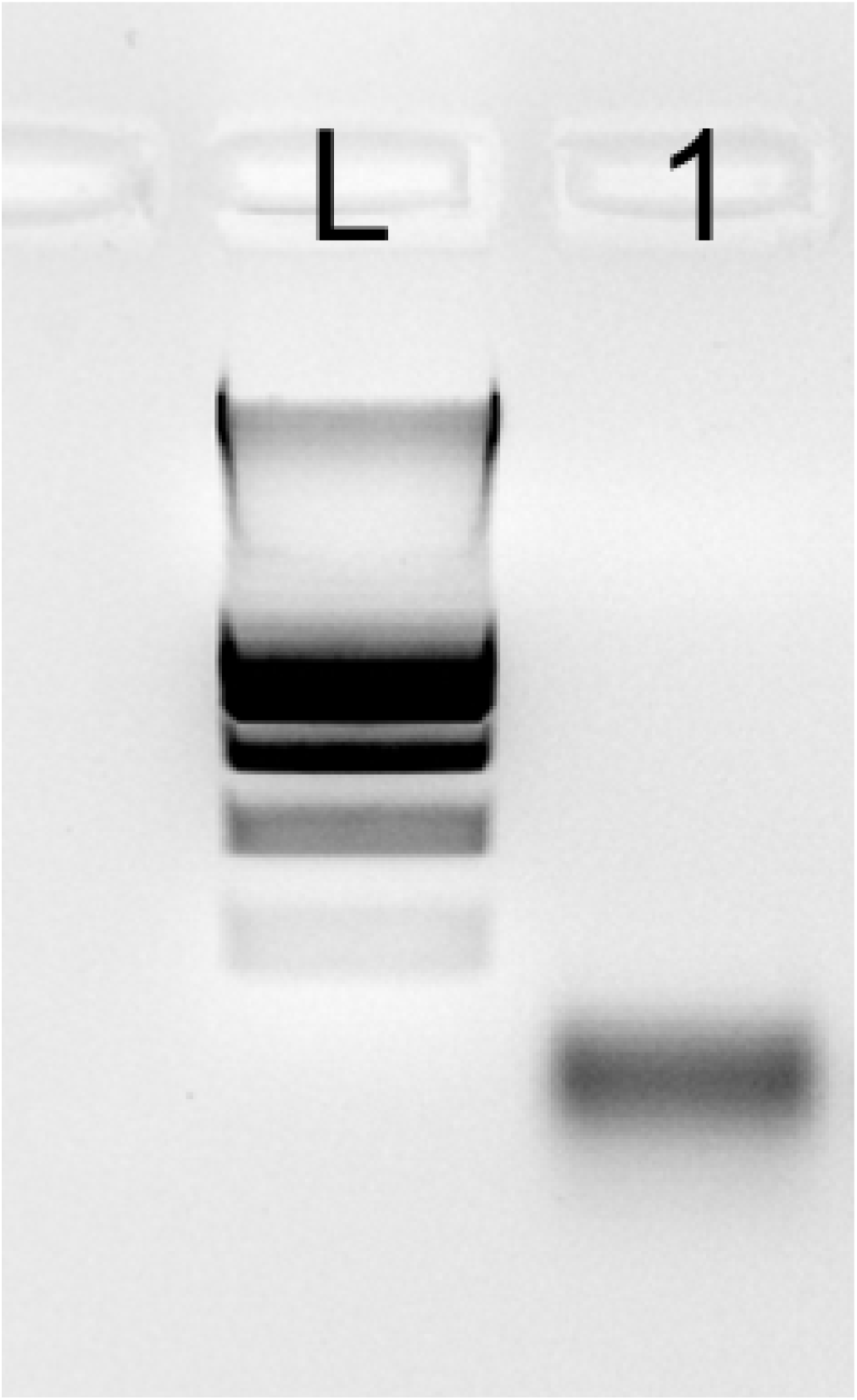
CasRx gRNA for tbxta gene.

**Additional file 9: Figure S4.**
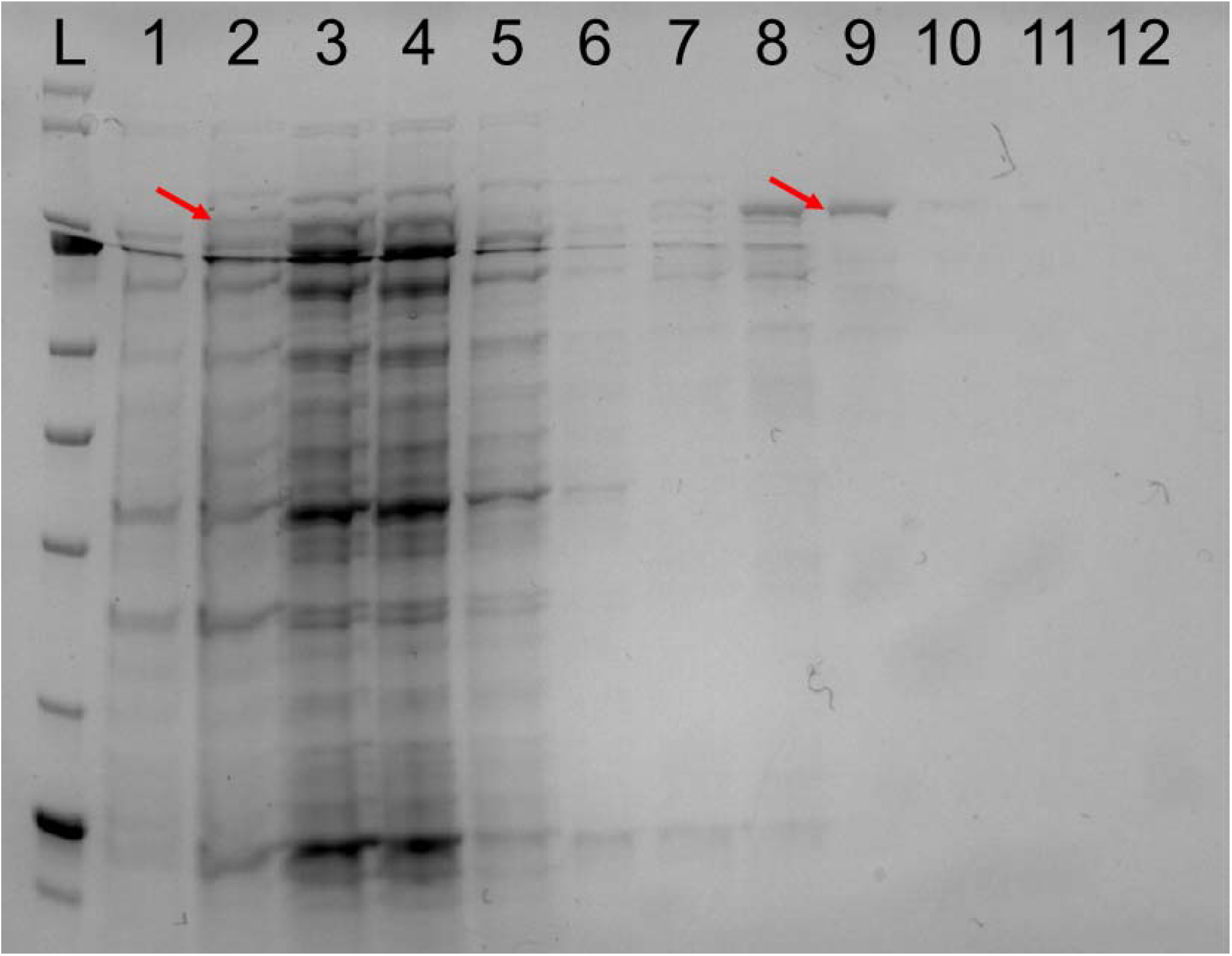
CasRx protein production.

**Additional file 10: Table S1.**
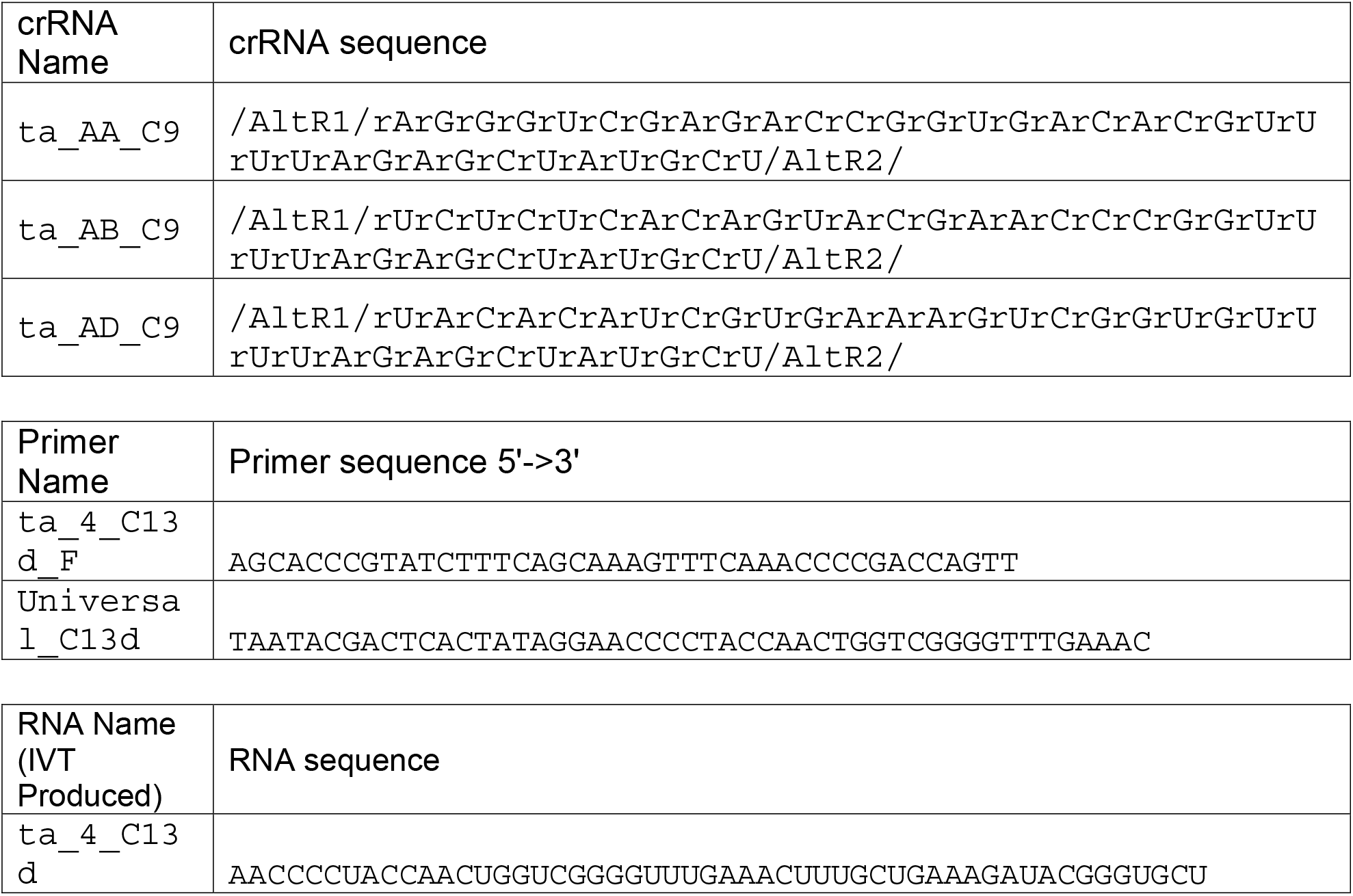
Primers and RNA sequences.

