## Supplementary figures and images for "Comparison of robotic automated and manual injection methods in zebrafish embryos for high throughput RNA silencing using CRISPR-CasRx"

### Figure S1

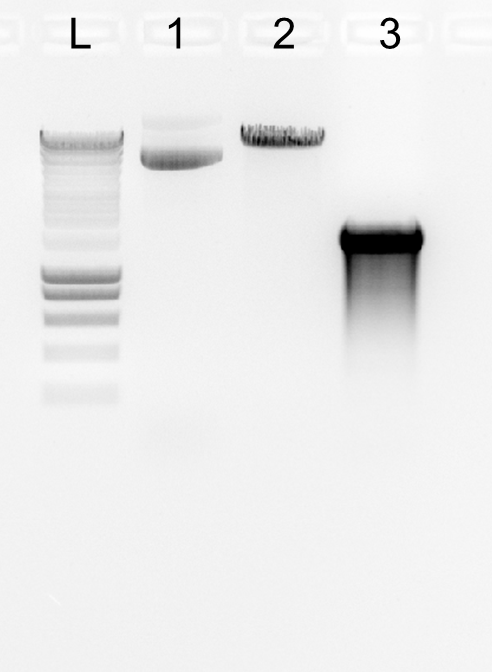

### Figure S2

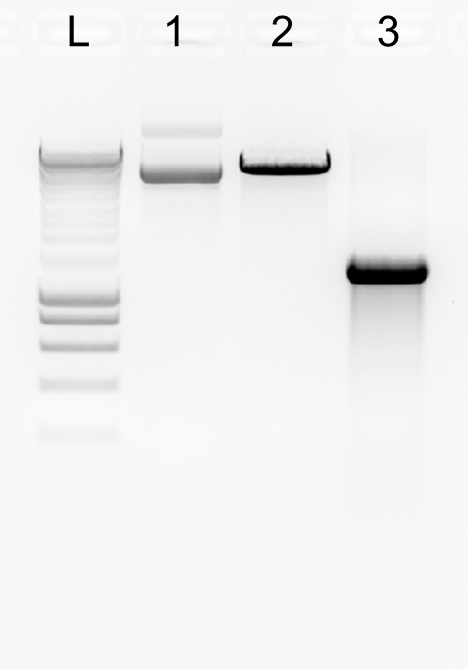

### Figure S3

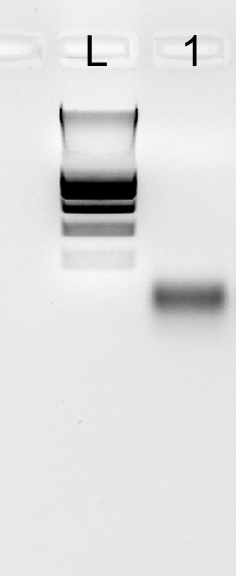

### Figure S4

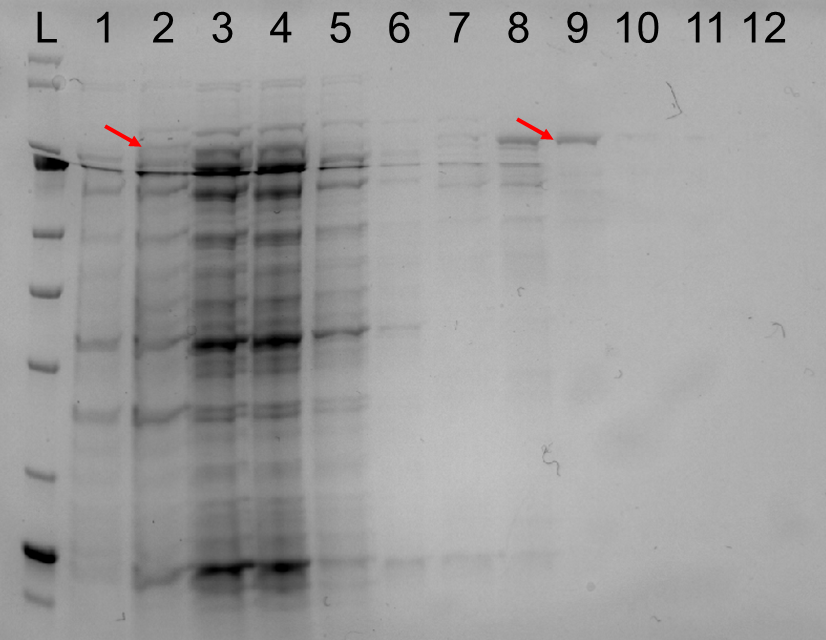
